# Single cell determination of cardiac microtissue structure and function using light sheet microscopy

**DOI:** 10.1101/2020.01.22.915926

**Authors:** Diwakar Turaga, Oriane B. Matthys, Tracy A. Hookway, David A. Joy, Meredith Calvert, Todd C. McDevitt

**Author notes:** Authors contributed equally (Diwakar Turaga, Oriane B. Matthys). **Corresponding Author:** Todd C. McDevitt, 1650 Owens St., San Francisco, CA 94158.

## Abstract

Native cardiac tissue is comprised of heterogeneous cell populations that work cooperatively for proper tissue function; thus, engineered tissue models have moved toward incorporating multiple cardiac cell types in an effort to recapitulate native multicellular composition and organization. Cardiac tissue models comprised of stem cell-derived cardiomyocytes require inclusion of non-myocytes to promote stable tissue formation, yet the specific contributions of the supporting non-myocyte population on the parenchymal cardiomyocytes and cardiac microtissues have yet to be fully dissected. This gap can be partly attributed to limitations in technologies able to accurately study the individual cellular structure and function that comprise intact 3D tissues. The ability to interrogate the cell-cell interactions in 3D tissue constructs has been restricted by conventional optical imaging techniques that fail to adequately penetrate multicellular microtissues with sufficient spatial resolution. Light sheet fluorescence microscopy overcomes these constraints to enable single cell-resolution structural and functional imaging of intact cardiac microtissues. Multicellular spatial distribution analysis of heterotypic cardiac cell populations revealed that cardiomyocytes and cardiac fibroblasts were randomly distributed throughout 3D microtissues. Furthermore, calcium imaging of live cardiac microtissues enabled single-cell detection of cardiomyocyte calcium activity, which showed that functional heterogeneity correlated with spatial location within the tissues. This study demonstrates that light sheet fluorescence microscopy can be utilized to determine single-cell spatial and functional interactions of multiple cell types within intact 3D engineered microtissues, thereby facilitating the determination of structure-function relationships at both tissue-level and single-cell resolution.

**Impact Statement:** The ability to achieve single-cell resolution by advanced 3D light imaging techniques enables exquisite new investigation of multicellular analyses in native and engineered tissues. In this study, light sheet fluorescence microscopy was used to define structure-function relationships of distinct cell types in engineered cardiac microtissues by determining heterotypic cell distributions and interactions throughout the tissues as well as by assessing regional differences in calcium handing functional properties at the individual cardiomyocyte level.

## Introduction

Engineered models of native tissue have become more physiologically accurate with increasing complexity and heterogeneity of cellular constituents. Engineered cardiac tissue depends on stromal cell contributions to enable robust tissue formation and stable culture as well as to promote cardiomyocyte phenotype and functional properties^1–10^. However, methods to dissect the multicellular organization and function of engineered tissue constructs have been typically limited to bulk tissue-level measures, as structural and functional analyses at single-cell resolution typically necessitate removing cells from their 3D environment prior to analysis. Technological advances with respect to single-cell measurements, specifically increased accessibility of single cell RNA sequencing technologies, have enabled transcriptomic analyses at the single cell level but lack the context of spatial information or functional outputs^11, 12^. Therefore, this study examined the ability of light sheet microscopy to interrogate structural and functional information of intact tissue constructs with single-cell resolution.

Analysis of 3D tissue structure typically requires 1) histological sectioning combined with epifluorescence microscopy, which allows for high resolution imaging of structure, but is limited by the need to physically slice the tissue, and 2) point-scanning fluorescence sectioning microscopes such as confocal or two-photon microscopes, which enable high spatial resolution imaging, but are hindered by speed limitations. Cardiac tissue function is typically assessed at the bulk tissue level (i.e. calcium imaging^4, 5, 7–9^ or contractile force testing^3, 7, 8, 13^ of entire microtissues) or at single cell resolution by dissociating tissues, culturing the cells in 2D for multiple days, and then performing patch clamp analysis^2, 4, 7^. In contrast, light sheet fluorescence microscopy (LSFM) allows for high resolution imaging of thick tissue samples at significantly faster speeds with low photobleaching and phototoxicity^14–16^. A cylindrical lens is used to create a sheet of light that illuminates only the section of the tissue in the focal plane of the objective lens, thus allowing for a high resolution camera to image the entire 2D plane simultaneously. LSFM has been used to image entire embryos with single cell resolution over multiple hours^14, 17^ and to image the function of hundreds of neurons simultaneously^15, 18–20^. This study analyzes multicellular organization of homotypic (cardiomyocytes alone) and heterotypic (cardiomyocytes and cardiac fibroblasts) microtissues by combining immunofluorescence staining and LSFM to dissect cell-type specific localization and calcium handling function of the engineered tissue constructs at high spatial and temporal fidelity. Coupling single-cell structural and functional information of 3D engineered tissues will advance understandings of cell type-specific contributions to tissue properties as well as enable further insights into developmental- or disease-specific biological events that can be interrogated in 3D tissue models.

## Materials and Methods

### Cardiac fibroblast cell culture

Human cardiac fibroblasts (CFs) were purchased from Cell Applications (lot #s 2584 & 3067; San Diego, CA) and cultured according to manufacturer’s recommendations: fibroblasts were seeded onto non-coated TCPS plates at density of 1×10^4^ cells/cm^2^ and cultured in Cardiac Fibroblast Medium (Cell Applications) for up to 10 passages. CFs were passaged by incubating with 0.25% Trypsin-EDTA for 5min when cultures reached ∼80% confluence.

### Cardiomyocyte differentiation

Human induced pluripotent stem cells (hiPSCs) (WTC11 cells modified with GCaMP6f reporter in the AAVS1 safe harbor locus^21, 22^; generously donated by Dr. Bruce Conklin) were seeded onto Matrigel-coated (80µg/mL; Corning, Corning, NY) plates at a concentration of 3×10^4^ cells/cm^2^ in mTeSR1 medium (Stem Cell Technologies, Vancouver, CA) supplemented with 10µM ROCK inhibitor (Y-27632, SelleckChem, Houston, TX) for the first 24h. Differentiation of hiPSCs to cardiomyocytes was performed using a serum-free, chemically defined protocol^23, 24^. Briefly, once hiPSCs reached 100% confluence (∼3-4 days; denoted as differentiation day 0), cells were fed with 12µM CHIR (SelleckChem) in RPMI1640 medium (Thermo Fisher, Waltham, MA) with B27 supplement without insulin (RPMI/B27-; Life Technologies, Grand Island, NY). After 24h, CHIR was removed by feeding with RPMI/B27-and on day 3, cells received a 48h-treatment with 5µM IWP2 (Tocris, Bristol, UK) in RPMI/B27-. Medium was then switched to RPMI1640 medium containing B27 supplement with insulin (RPMI/B27+; Life Technologies) and fed every 3 days thereafter. On day 15 of differentiation, hiPSC-CMs were re-plated onto Matrigel-coated plates at a density of 1×10^5^ cells/cm^2^ in RPMI/B27+ with 10µM ROCK inhibitor. Selection of CMs was achieved by lactate purification with two 2-day feedings with no-glucose Dulbecco’s Modified Eagle Medium (Thermo Fisher) supplemented with 1X Non Essential Amino Acids (NEAA; Corning), 1X Glutamax (*L*-glut; Life Technologies), and 4mM Lactate)^25^. After lactate selection, cultures were returned to RPMI/B27+ media and re-fed every 3 days thereafter with fresh media.

### Cardiac microtissue formation

Lactate-purified hiPSC-CMs and primary human CFs were dissociated with 0.25% Trypsin for 10min to obtain a single-cell suspension, mixed at a 3:1 CM:CF ratio, and seeded into an array of inverted 400µm pyramidal microwells at a density of ∼2000 cells per microwell^9,26^. Cells self-assembled into 3D tissues over the course of 24h and were then transferred from the microwells to rotary orbital suspension culture at a density of ∼4000 microtissues per 10cm Petri dish (∼8×10^5^ cells/mL) and maintained in RPMI/B27+ medium^26^ until analysis.

### Immunofluorescence staining

Microtissues were fixed in 10% neutral-buffered formalin for 1h at RT and then washed 3x with PBS. Samples were permeabilized in 1.5% Triton X-100 (Sigma-Aldrich, St. Louis, MO) for 1h and blocked overnight at 4°C in 2% normal donkey serum and 0.1% Tween-20. Tissues were incubated in primary antibody against GATA4 (1:50 dilution; Santa Cruz Biotechnology, Dallas, TX) overnight at 4°C, and counterstained with Alexa Fluor 555 (1:400; Thermo Fisher) and Hoechst (1:1000; Thermo Fisher) overnight at 4°C (Supplementary Table 1 for antibody information).

### Structural light sheet microscopy

Cardiac microtissues stained for GATA4 and labeled with Hoechst were suspended in size 2 glass capillaries (Zeiss; ∼1mm inner diameter) in 2% low-melt agarose (made up in PBS; IBI Scientific, Dubuque, IA) immediately prior to imaging (Supplementary Figure 1). The Zeiss z.1 light sheet microscope used for imaging was equipped with two PCO.edge sCMOS cameras, 10x 0.2 NA illumination lens, 20x 1.0 NA detection lens, and 488/647nm lasers for dual imaging. Cardiac microtissue samples (n ≥ 9 per condition) were each imaged at three angles (120° rotations between views), and then stitched with multi-view reconstruction to provide isotropic resolution throughout the microtissue. Volumetric reconstruction and size analyses of the microtissues were performed using custom Matlab (R2019a) scripts (adapted from^15^).

### Cell classification and spatial quantification

Imaris image analysis software (version 9.3.1) was used to identify labeled cell nuclei within the microtissues. Classification of cell identity was performed by determining co-localization of DAPI+ and GATA4+ nuclei. CMs were identified as GATA4+ nuclei whereas CFs were classified as cells with GATA4-nuclei. The spatial coordinates for each CM and CF were used to determine the multicellular arrangement within the microtissues. The numbers of nearest CM or CF neighbors (within a 20μm radius) for each CM were calculated using a custom python script to create 3D spatial maps of CM homotypic and heterotypic interactions. The nearest-neighbor calculation was performed across multiple CM+CF microtissues (n=8) to determine the distribution of proximal interactions. This empirical distribution was compared against a simulated distribution of randomly-dispersed CMs and CFs. CMs and CFs were simulated as 10µm-diameter hard spheres randomly dispersed inside a larger spherical volume, matching the empirical parameters of microtissue size (average CM+CF microtissue diameter of 165µm; Figure 1) and multicellular composition (average of 400 CMs and 127 CFs per heterotypic microtissue; Supplementary Figure 2). The simulation was performed using a custom python script to generate a random sequential packing of hard spheres for the total number of cells within a volume (517), followed by proportional random assignment of cell identity as either CM (400/517) or CF (127/517) to each sphere^27^.

**Figure 1.**
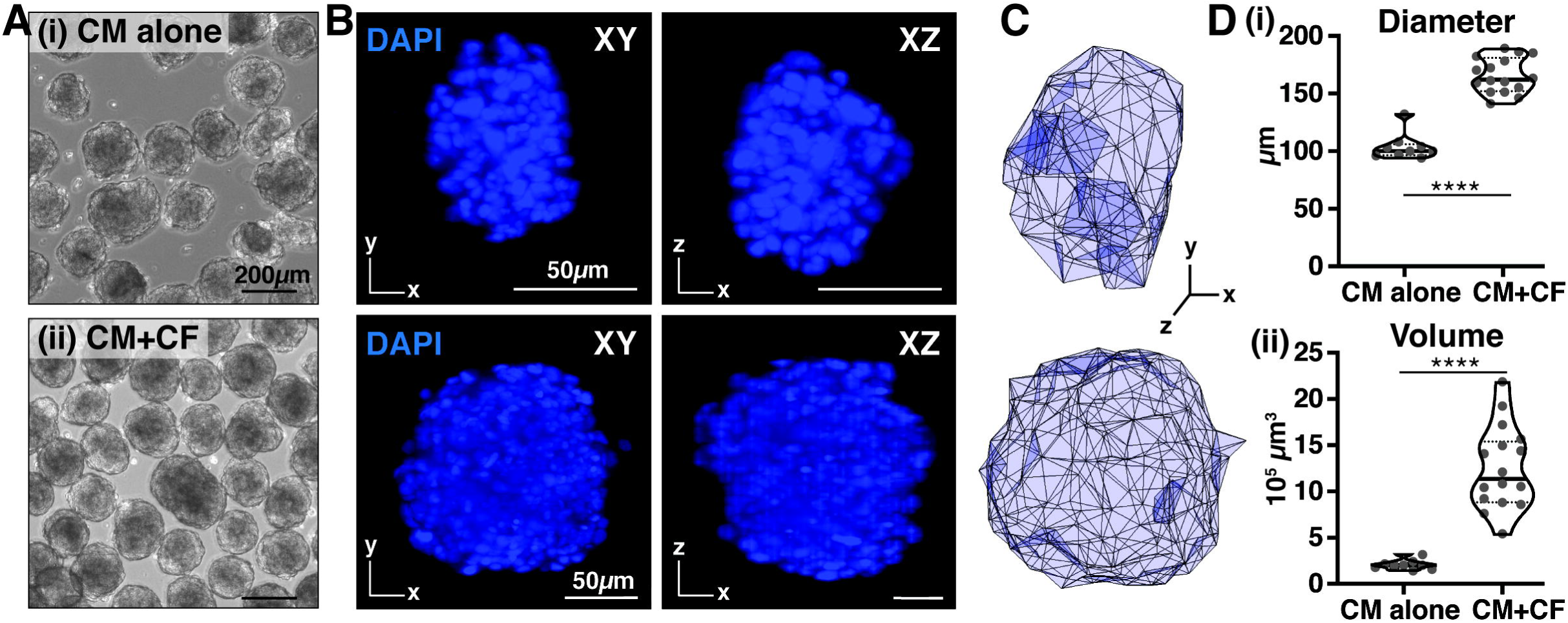
Multi-view 3D imaging of engineered cardiac microtissues. (A) Phase images of microtissues comprised of only cardiomyocytes (i) or cardiomyocytes with cardiac fibroblasts (ii). Scale bar = 200µm. (B) XY and XZ images obtained from multi-view imaging (120° rotation) of CM alone and CM+CF microtissues. Scale bar = 50µm. (C) 3D volumetric reconstruction of cardiac microtissues using Imaris software. (D) Size measurements of reconstructed cardiac microtissues. Cardiac microtissues containing CFs were larger in diameter and volume compared to tissues comprised of only CMs. **** p<0.0001 by unpaired t-test with Welch’s correction. The variance of standard deviations between CM alone and CM+CF microtissue was not significant for tissue diameter, but was significant for tissue volume (p < 0.0001 by the Brown-Forsythe test).

### Functional light sheet microscopy

5-10 live cardiac microtissues were suspended in size 2 glass capillaries in 2% low-melt agarose made up in Tyrode’s solution (137mM NaCl, 2.7mM KCl, 1mM MgCl_2_, 0.2mM Na_2_HPO_4_, 12mM NaHCO_3_, 5.5mM D-glucose, 1.8mM CaCl_2_; Sigma-Aldrich). Calcium handling properties were assessed in live cardiac microtissues by recording fluorescence intensity of GCaMP6f calcium indicator while tissues were maintained at 37°C. 3D calcium imaging stacks were imaged at ∼120ms per frame and each z-spacing was 0.86μm per frame. Single optical sections (3-4μm light light thickness) were excited at 488nm and imaged at ∼20Hz for 52 seconds (1000 total frames). Regions of interest (ROIs) were manually selected for individual CMs, and normalized fluorescence intensity change (Δ F/F) profiles were calculated for each calcium transient. A custom Matlab script was used to perform K-means clustering (k=2) on the calcium transient profiles to determine synchronously-active CM calcium profiles.

### Statistics

The mean and standard deviations for cardiac microtissue size analyses (Figure 1) were calculated for independent CM alone (n = 9) and CM+CF (n = 16) tissues. Statistical testing of diameter and volume means was performed using unpaired t-test with Welch’s correction and variance of diameter and volume standard deviations was analyzed using the Brown-Forsythe test. A two-sided Kolmogorov-Smirnov (K-S) statistical test was performed to determine significance between the empirical vs. simulated distributions of homotypic and heterotypic nearest neighbors (Figure 2E). The Brown-Forsythe statistical test was used to determine significance between the variance of inter-beat intervals between CM alone and CM+CF microtissues (Figure 3). All statistical tests were performed using SciPy^28^ (version 1.3.1) with significance determined at p < 0.05.

**Figure 2.**
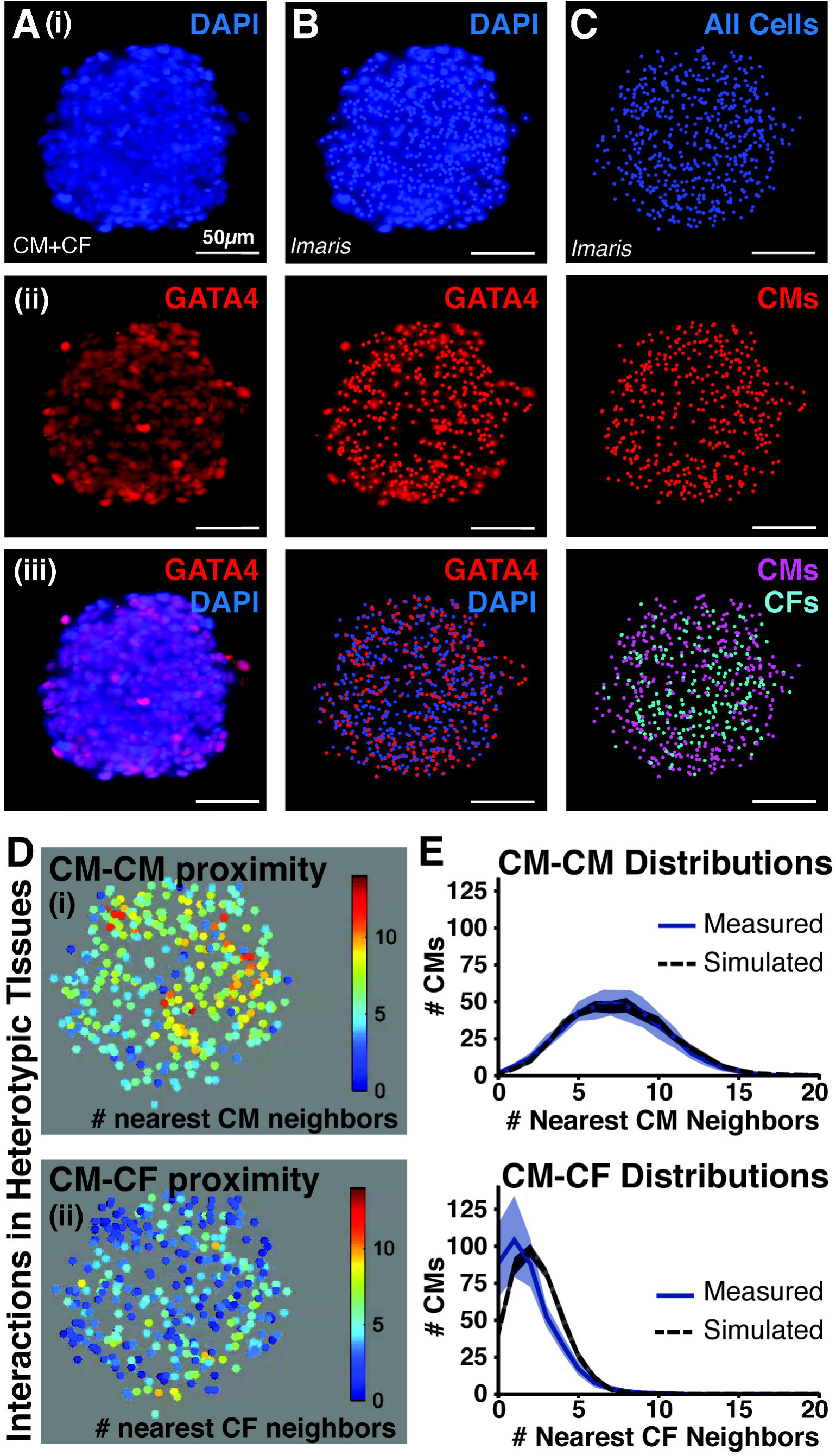
*In situ* cell classification enables cell-specific spatial quantification of heterotypic CM+CF microtissues. (A) Maximum intensity projections of multi-view images were obtained for microtissues with labeled nuclei (DAPI; i), GATA4 staining (ii), and merged channels (iii). Scale bar = 50µm. (B) Nuclear localization within maximum intensity projection multi-view images. (C) Cell classification based on nuclear localization of DAPI (marking all cells) and GATA4 (identifying CMs). GATA4+ nuclei were classified as CMs whereas GATA4-nuclei were classified as CFs. (D) Number of nearest homotypic (CM-CM; i) and heterotypic (CM-CF; ii) neighbors for each CM in the heterotypic CM+CF microtissue. (E) Measured distributions of numbers of CM (i) and CF (ii) nearest neighbors matched that of a simulated tissue model of randomly-distributed CMs and CFs, indicating that CFs were randomly distributed throughout CMs in the empirical microtissues.

**Figure 3.**
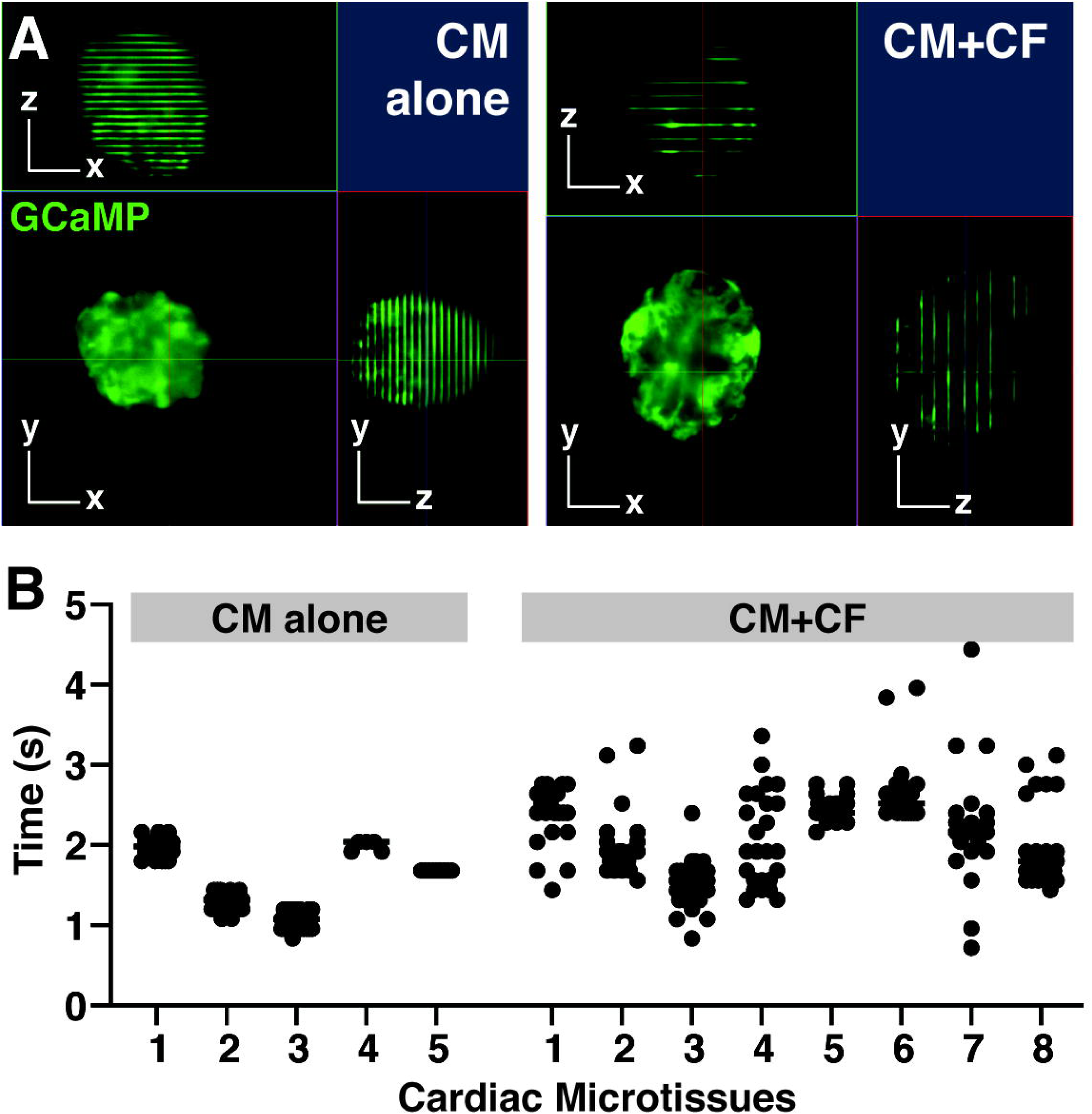
Light sheet calcium imaging of live 3D cardiac microtissues. (A) Representative orthogonal (XY, XZ, YZ) images obtained from z-stack scans through spontaneously beating CM alone and CM+CF microtissues. Fluorescent lines represent the calcium flux of the tissue beat during acquisition (acquisition parameters: ∼120ms per frame, 0.86μm spacing per frame). Scale bar = 50µm. (B) Time between spontaneous beats (inter-beat interval) of each microtissue calculated based on z-scanned images. Microtissues containing CFs displayed higher variability of inter-beat intervals compared to CM alone microtissues (p < 0.0001 by the Brown-Forsythe test).

## Results

### Multi-view light sheet imaging enables high resolution 3D reconstruction of cardiac microtissues

Cardiac microtissues comprised of either hiPSC-cardiomyocytes alone (CM alone; Figure 1A(i)) or hiPSC-CMs with cardiac fibroblasts (CM+CF; Figure 1A(ii)) were imaged at multiple views (120° apart; lateral resolution = 0.5μm × 0.5μm; axial resolution = 3-4μm) in order to obtain improved and consistent resolution across the entire microtissue (Figure 1B). The resultant higher resolution imaging enabled accurate 3D volumetric renderings of individual microtissues (Figure 1C; Supplementary Movie 1), which permitted calculation of tissue size and shape. Despite seeding the same number of total cells for both homotypic and heterotypic tissues, cardiac microtissues that contained CFs were ∼60% larger in diameter and ∼6-fold greater in volume than the CM alone microtissues (average diameter of 165µm vs. 103µm with p-value = 1.80×10^-10^; average volume of 12.5×10^5^µm^3^ vs 2.1×10^5^µm^3^ with p-value = 1.08×10^-7^; Figure 1D), though CM alone cardiac microtissues were more consistent in their size as compared to CM+CF microtissues (respective volume standard deviations of ±0.5×10^5^µm^3^ compared to ±4.3 x10^5^µm^3^; p-value = 0.002 by Brown-Forsythe test of volume variance, with statistical significance determined at p = 0.05). Identification of DAPI-labeled cells by Imaris image analysis revealed an average of 165 cells in the CM alone microtissues compared to 527 cells in the heterotypic CM+CF microtissues (Supplementary Figure 2).

### Localization of labeled cells allows for cell identity classification and intercellular spatial analyses

Multi-view imaging for different fluorescent channels was performed to identify CMs (GATA4+ nuclei; Figure 2A(ii)) from all cells (GATA4-nuclei; Figure 2A(i)) and individual cells were segmented using Imaris image analysis software to identify the *in situ* location of individual cells (Figure 2B). Therefore, the classification of cell identity with respect to 3D spatial location within the microtissue was determined by combining localization information for the different cell types. CMs were identified as cells with GATA4+ nuclei while CFs were classified as cells with GATA4-nuclei (Figure 2C). Furthermore, counts of classified CMs and CFs across analyzed heterotypic cardiac microtissues revealed that the seeding ratio of 3:1 CMs:CFs was maintained through culture and image analysis (Supplementary Figure 2). Taken together, the image analysis pipeline of multi-view reconstruction, cell localization, and identity classification resulted in 3D spatial mapping of CMs and CFs in each microtissue (Supplementary Movie 2).

The interrogation of heterogeneous cellular packing within cardiac microtissues could be derived from the 3D spatial mapping of CMs and CFs. Heterotypic (CM+CF) cardiac microtissues were utilized in order to study intercellular interactions between different pairings of cell types (i.e. homotypic (CM-CM) versus heterotypic (CM-CF) interactions). The local density of homotypic and heterotypic neighbors was determined for each individual CM (Figure 2D). On average, each CM was located adjacent to 6-8 CMs and 1-2 CFs within a 20µm radius. Furthermore, looking at the spatial distribution of the interactions across 3D microtissues revealed that homotypic interactions (yellow-to-red heatmap range; Figure 2D(i)) were greater in the center of the tissue compared to the edge, while heterotypic CM-CF interactions were generally consistent throughout the microtissues (blue-green heatmap range; Figure 2D(ii)). To assess the extent to which CMs and CFs were distributed in a random or biased manner throughout the microtissues, the empirical distributions of nearest homotypic and heterotypic neighbors were compared to a simulated model of well-mixed, randomly-dispersed heterotypic tissues that matched empirical tissue size and cellular composition (Figure 2E). The simulated distribution curves indicated that CMs should be surrounded by 6-8 CMs and 2-3 CFs on average. The empirical and simulated nearest-neighbor distributions were not significantly different as determined by two-sided K-S test, indicating that the CFs in the imaged heterotypic microtissues were randomly dispersed among the CMs. The K-S value for homotypic (CM-CM) distribution analysis was 0.0264 and the K-S value for heterotypic (CM-CF) distribution analysis was 0.1630, where K-S > 1.224 indicates a statistically significant difference between the two distributions (p < 0.05). Furthermore, radial distribution of cells throughout the empirical imaged tissues did not differ significantly from the simulated tissue model (K-S = 0.2386; Supplementary Figure 3). Therefore, Imaris analysis detected cells at the microtissue center just as well as at the tissue edge, indicating that the accuracy of cell detection did not diminish despite attenuation of imaging resolution with increasing tissue depth.

### Live light sheet calcium imaging enables detection of functional heterogeneity between engineered cardiac microtissues

In order to study functional synchrony within individual tissues as a result of multicellular composition, calcium imaging of live cardiac microtissues was performed. Microtissues comprised of hiPSC-CMs expressing a genetically-encoded calcium indicator, GCaMP6f, enabled direct visualization of synchronicity of calcium handling activity throughout tissues, as well as individual CM calcium fluxes within single optical sections. The periodicity of spontaneous calcium propagation was determined by z-stack imaging through cardiac microtissues (Supplementary Movie 3). Combining the known z-scan rate with the measured distance between beats (fluorescent lines indicating GCaMP6f signal) in the orthogonal (XZ/YX) views of the image (Figure 3A) enabled the determination of inter-beat time interval for each microtissue (Figure 3B). Microtissues comprised of only CMs beat more periodically than the microtissues containing CFs, as exhibited by the smaller variation in the CM alone inter-beat intervals compared to the more widespread values of the CM+CF tissues (inter-beat interval standard deviation of ±0.095ms for CM alone microtissues and ±0.447ms for CM+CF tissues; p-value = 6.865×10^-9^ by Brown-Forsythe test of inter-beat interval variance, with statistical significance determined at p = 0.05). The orthogonal views of z-stack calcium activity displayed distinct lines of GCaMP6f fluorescence that transected the entirety of the CM alone microtissue (Figure 3A), suggesting that CMs within the single z-plane were typically firing in synchrony and therefore the variation in inter-beat interval periodicity was due to time rather than space. The GCaMP6f lines transecting the CM+CF microtissues exhibited some breaks in the fluorescence, likely indicating the presence of CFs in those particular locations.

In order to quantitatively determine the synchronicity of calcium transients of individual CMs within the microtissues, a time series of a single optical section within the microtissues was captured at ∼20Hz. The two CM+CF microtissues in the same optical field of view beat spontaneously but independently from one another (Supplementary Movie 4). ROIs for individual CMs were selected in both tissues (Figure 4A) and the normalized fluorescence intensity traces of each ROI were plotted; calcium traces from the top tissue are depicted in red and the lower tissue traces are depicted in blue (Figure 4B, Supplementary Figure 4). Unbiased k-means clustering grouped CMs with similar calcium transients. The clustered calcium traces partitioned entirely with respect to the microtissues they originated from, indicating that CMs from the two tissues fired at independent times and rates from one other (Figure 4C). However, although CMs within the individual tissues fired synchronously, differences in calcium transient duration varied, even among CMs in close proximity to one another (Figure 4C’, arrows) sustained longer calcium traces than their neighbors. Therefore, this analysis platform demonstrates that regional analysis of individual CM calcium transients can be used to assess functional heterogeneity as it relates to spatial location.

**Figure 4.**
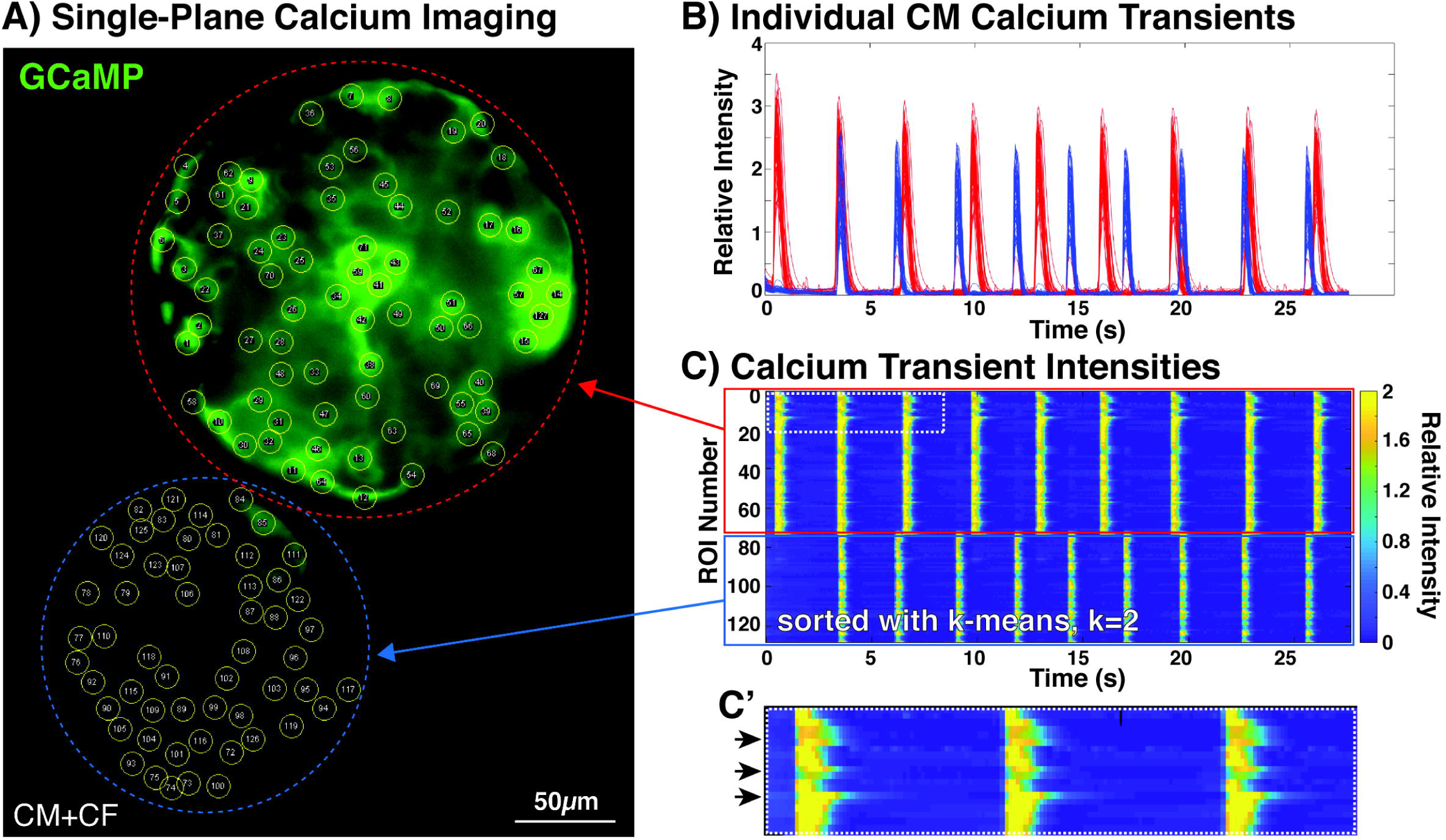
Single plane calcium imaging of live cardiac microtissues. (A) Single optical section of two cardiac microtissues with regions of interest (ROIs) selected over individual CMs. Scale bar = 50µm. (B) Normalized calcium transient traces for each ROI (n = 126). (C) K-means clustering of ROI normalized fluorescence activity enables determination of functional heterogeneity. (C’) Inset of ROI clustering highlighted differences in calcium transient duration.

## Discussion

This study interrogated the structural and functional information of intact 3D engineered cardiac microtissues at single-cell resolution. The methods described establish a powerful toolkit to better dissect *in situ* multicellular heterogeneity and the impacts of organization on function. Light sheet fluorescence microscopy was used to map 3D tissue structure by segmenting, localizing, and classifying distinct cardiac cell populations. Use of LSFM also enabled imaging of multiple live cardiac tissues in a rapid manner to assess functional tissue synchrony as well as detect individual cell functional variation at different depths and spatial locations within intact engineered tissues.

Constructing a tissue-level representation from cells imaged at single-cell resolution provided an accurate model of engineered microtissue surface terrain and volumetric shape information (Figure 1). Although spatial resolution was greatest at the exterior edges of the tissues and gradually declined towards the center (Supplementary Movie 1), single-plane imaging at different depths of individual tissues confirmed the presence of cells distributed throughout the interior. The addition of clearing and refractive index-matching processes could improve the attenuation of resolution with depth of imaging^29–31^. However, despite decreased spatial resolution at the center of imaged microtissues, detection of cells by computational segmentation did not change based on tissue radius, thereby highlighting the ability of LSFM to accurately capture 3D multicellular density *in situ* (Supplementary Figure 3).

3D volumetric reconstruction of cardiac microtissues allowed for tissue-scale size analyses. Conventional methods to quantify tissue size include 2D cross-sectional measures of tissue slices as well as standard light microscopy image analysis tools, but these approaches are limited to measures along one plane. 3D reconstructions from the imaged cellular constituents therefore retain more accurate size and shape information. The heterotypic CM+CF cardiac microtissues were larger than the homotypic CM microtissues despite identical initial seeding conditions (Figure 1D), suggesting that CFs impact tissue formation and culture. In the native heart, CFs interact with CMs directly via physical adhesion molecules and indirectly via secretion and organization of the surrounding extracellular matrix (ECM)^32–36^. These CF-mediated methods of intercellular interactions could potentially account for the larger heterotypic microtissue size distinction by more strongly promoting the adhesion of cells in the initial tissue formation phase, ultimately leading to larger numbers of cells assembling into the microtissue constructs. The distinct tissue-level structural differences between microtissue compositions was further analyzed at single-cell resolution by quantifying multicellular organization within the microtissues, in order to determine whether the addition of a stromal population changed the CM interaction properties.

Identification of cell number and identity was performed on the heterotypic (CM+CF) microtissues to compare to the initial seeding conditions (Figure 2). Although CMs and CFs were mixed at a 3:1 ratio and seeded at a total of 2000 cells per tissue, an average of only ∼500 cells were identified in the heterotypic microtissues, but the ratio of CMs to CFs was retained at an average of 400 CMs to 127 CFs (∼3:1 CM:CF) (Supplementary Figure 2). The lower-than-expected cell numbers were likely due to lack of total incorporation of all 2000 cells during the initial tissue formation step—which is to be expected^9,26, 37^. However, the maintained ratio of the heterogeneous cardiac cell populations allowed for the interrogation of intra-tissue spatial interactions. Cell identity was classified based on positive or negative staining for GATA4, a nuclear cardiac marker, along with DAPI labeling of individual nuclei in order to identify CMs (Figure 2A-C). Specific phenotypic markers for CFs are particularly challenging^38, 39^, therefore, a subtractive method was used to distinguish CFs from CMs in the microtissues. Alternatively, constitutive expression of a fluorescent protein could be used to label non-myocytes prior to tissue formation, thereby improving longitudinal analyses of multicellular, heterotypic interactions. Based on cell identity classification and spatial localization of cells within heterotypic microtissues, CFs appeared to be randomly distributed among the CMs (Figure 2D,E). Furthermore, random-dispersion simulations confirmed that pockets slightly enriched for heterotypic (CM-CF) interactions arise spontaneously in randomly mixed cardiac microtissues, suggesting a mechanism for multicellular CF organization driven by stochastic inhomogeneity in the initial cell mixture. This multicellular spatial analysis is important as it can be used to determine cell-specific localization biases within complex organizational tissue structures.

The ability to dissect structure-function-phenotype relationships at the single-cell level within engineered tissue constructs would advance understanding of how multicellular spatial arrangements impact functional heterogeneity. With increased access to and improved robustness of single cell RNA sequencing technologies, studying tissue transcriptional phenotypes at the single cell level has become increasingly widespread^11, 12^, yet most assessments of tissue function are still analyzed at the bulk tissue-level. This study, however, used LSFM to describe novel methods for imaging live cardiac tissue functional properties at single-cell resolution. Since GCaMP6f was used to visualize calcium transients, no exogenous dye was needed for imaging, though this method is compatible with the use of fluorescent calcium and action potential dyes. In order to assess tissue-level functional synchrony before focusing in on individual cell activity, calcium imaging of live cardiac microtissues was first acquired by scanning through the microtissue at a fixed z-stack rate. The orthogonal imaging views (XZ, YZ) displayed fluorescent lines when the tissues beat during the z-stack acquisition (Figure 3A), which allowed the periodicity of beat rate to be determined. While the individual tissues had different intrinsic average beat rates, the variance of inter-beat intervals (time between sequential beats) was lower in the CM alone microtissues compared to the inter-beat interval variance of CM+CF microtissues (Figure 3B). Since cardiac tissue calcium dynamics are too fast to acquire full volumetric renderings via z-scanning, time series acquisition of single optical sections acquired at multiple z-positions throughout the tissue was needed to determine 3D connectivity network of calcium activity. Fixed plane calcium imaging also allowed for the assessment of calcium handling function at single-cell resolution within the engineered cardiac microtissues.

Although the CM calcium activity between independent tissues differed with respect to spontaneous beat rate, the calcium transients of individual CMs were largely synchronous within any single cross-section of heterotypic cardiac microtissues. (Figure 4). Unbiased k-means clustering of individual CM calcium transients identified cells that behaved most similarly to one another (Figure 4C). CM function was linked to spatial location by correlating the grouped calcium transients back to the specific CMs from which the traces were derived. Therefore, this method is uniquely poised to answer questions related to spatially-distinct functional heterogeneity within engineered constructs. For example, a pacemaker-like cell population within cardiac tissue could be detected by identifying the cells that originate calcium or action potential propagations. The ability to dissect the heterogeneity of cellular structure-function would be a powerful tool for the study of certain cardiac diseases. For example, conductive disorders, such as long QT syndrome (LQTS) and catecholaminergic polymorphic ventricular tachycardia (CPVT), induce abnormal heart rhythms due to ion channelopathies^40^. Spatial mapping of 3D engineered tissue models of LQTS or CPVT, created from primary and/or iPSC-derived cells, could be used to examine the mechanisms responsible for the dysregulation of action potential and calcium transient propagation associated with the pathology.

Current limitations in the functional imaging acquisition pipeline preclude 3D calcium imaging of the entire microtissues at single-cell resolution. While single planes within tissues can be imaged at 20Hz, it takes several seconds to image an entire z-stack at the fastest acquisition rate. This acquisition speed limitation could potentially be overcome by acquiring high-speed single plane time series every few micrometers apart within the tissue followed by post-imaging stitching of these frames, ultimately constructing a 3D network of functional propagation—a method known as post acquisition synchronization^41^. Another challenge is that additional in-depth quantitative assessments of calcium handling properties could not be determined using the Zeiss z.1 microscope. The calcium response of cardiac tissues to electrical stimulation at increasing frequencies reveals the relative maturity level of cardiomyocyte contractile machinery. However, electrical stimulation could not be performed due to steric constraints of the metallic chamber used in the light sheet microscope; therefore, using a non-metallic chamber could potentially circumvent this limitation. Other methods of stimulation, such as optogenetic^42, 43^ as opposed to voltage- or current-driven, could also be incorporated to control CM contractility and eliminate intrinsic differences in beat rate between independent tissues.

Altogether, this study demonstrates the ability to interrogate the structural and functional properties of intact, dense, 3D microtissues at single-cell resolution. Analogous to how single-cell RNA sequencing has led to significant advances in understanding the phenotypic heterogeneity within complex multicellular environments, single-cell imaging enabled by LSFM will improve the coupling of spatial organization and functional heterogeneity within engineered tissues. Ultimately, the convergence of parallel advances in complementary single-cell technologies will lead to a more comprehensive view of the individual contributions of heterotypic cells to integrated properties at the tissue-level.

## Acknowledgements

The authors acknowledge funding support from the California Institute of Regenerative Medicine (LA1-08015) and the Gladstone BioFulcrum Heart Failure Research Program. D.T. was supported by the Eli and Edythe Broad Regenerative Medicine and Stem Cell Fellowship (7000-136209-7028606-41-FELOW). O.B.M. is a National Science Foundation Graduate Research Fellow (1650113). T.A.H. was supported by an American Heart Association Postdoctoral Fellowship (15POST22750003). The authors would like to thank the Gladstone Histology and Light Microscopy Core and the Gladstone Stem Cell Core (Roddenberry Stem Cell Foundation). The authors also thank Dr. Nathaniel Huebsch and Dr. Bruce Conklin for providing the WTC11-GCaMP6f hiPSC line.

---

**Supplementary Figure 1. Microtissue sample preparation for light sheet microscopy.** Live or fixed cardiac microtissues (A) were placed into a microcentrifuge tube and allowed to settle. The supernatant liquid was aspirated and 1.5% low-melt agarose (microwaved then cooled to 37°C) was added to the microtissues in the tube. A glass capillary with plunger was lowered into the warmed agarose-microtissue suspension and the plunger was slowly raised to draw up the microtissues into the capillary (B). The capillary with the loaded microtissues was cooled at RT until the agarose solidified (C). Once the capillary was mounted in the light sheet microscope, the plunger was pushed down to extrude the agarose/microtissues from the capillary into the imaging field of the objectives.

**Supplementary Figure 2. Cell counts identified from 3D Imaris-reconstructed images.** An average of 165 cells were identified in homotypic (CM alone) microtissues and an average of 527 cells were found in heterotypic (CM+CF) microtissues, based upon DAPI labelling (top). Classification of cardiomyocytes (CMs) and cardiac fibroblasts (CFs) in heterotypic cardiac microtissues based upon GATA4 staining identified averages of 400 CMs and 127 CFs, resulting in an average ratio of 3.1:1 CM:CF (bottom). n = 9 CM alone microtissues. n = 16 CM+CF microtissues.

**Supplementary Figure 3.** Distribution of cell counts as a function of tissue radius for empirical heterotypic cardiac microtissues and simulated heterotypic tissues.

**Supplementary Figure 4. Normalization of calcium transient profiles for individual CMs.** (A) ROI1 and ROI2 were selected around different CMs from the top microtissue. (B) Raw traces of fluorescence intensity. (C) Calcium transients were normalized by calculating the change in fluorescence intensity divided by the baseline intensity (ΔF/F)

**Supplementary Movie 1.** 3D scan through a multi-view reconstructed GATA4-stained cardiac microtissue.

**Supplementary Movie 2.** Cell localization and classification of heterotypic cardiac microtissues using Imaris image analysis software.

**Supplementary Movie 3.** 3D z-scan imaging of live cardiac microtissue calcium activity.

**Supplementary Movie 4.** Single-plane calcium imaging of live heterotypic cardiac microtissues.

