## Supplementary figures and images for "Single cell determination of cardiac microtissue structure and function using light sheet microscopy"

### Supplemental Figure 1

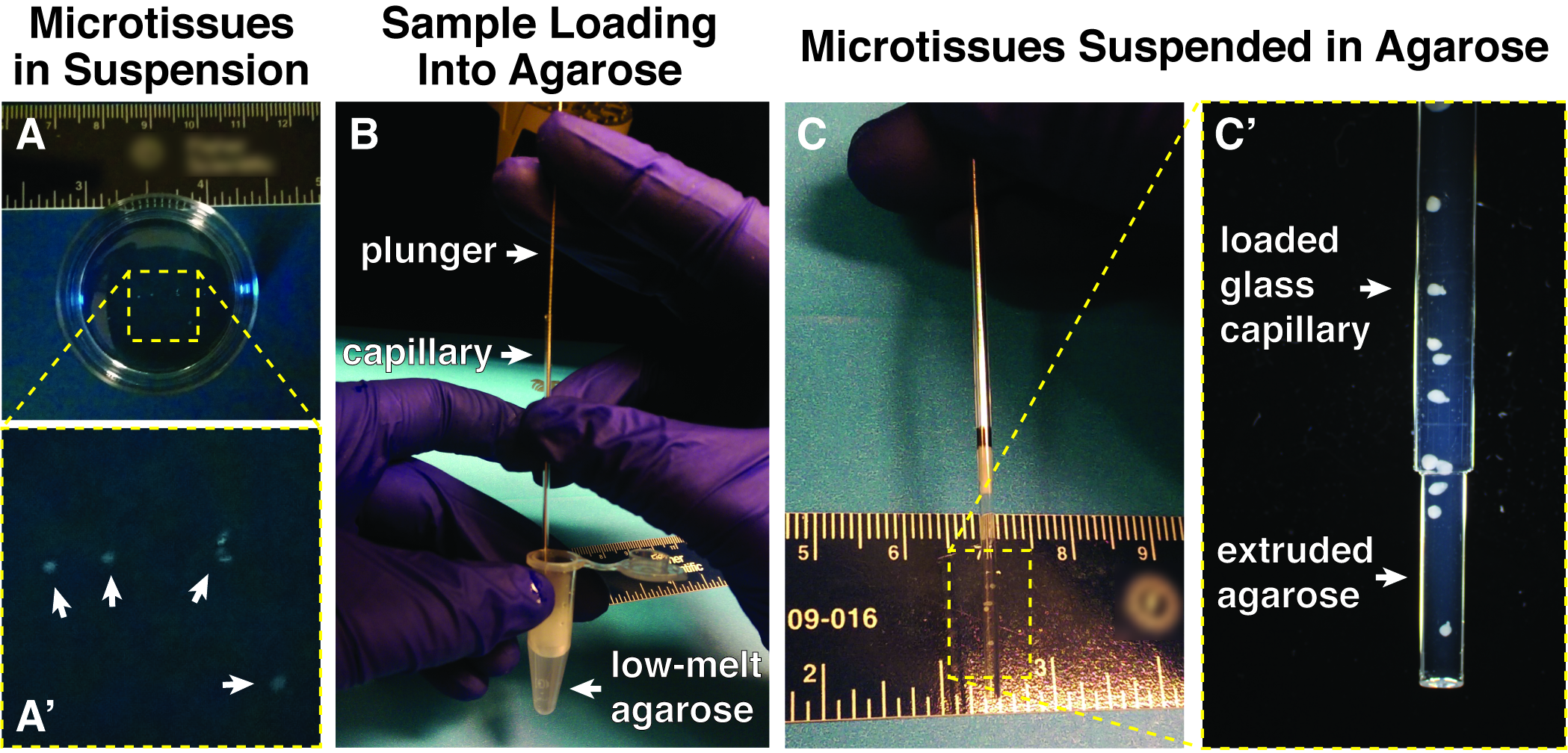

### Supplemental Figure 2

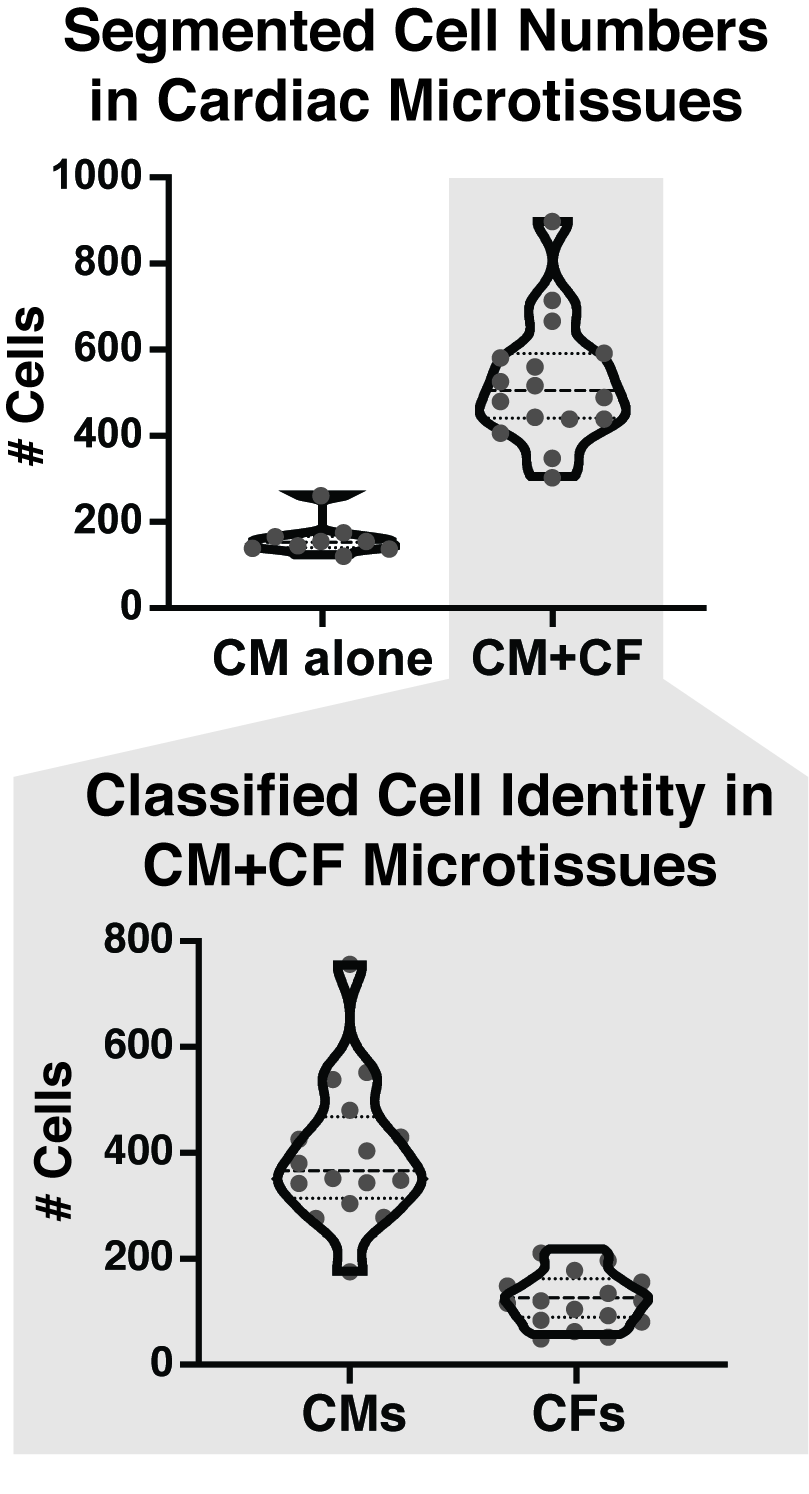

### Supplemental Figure 3

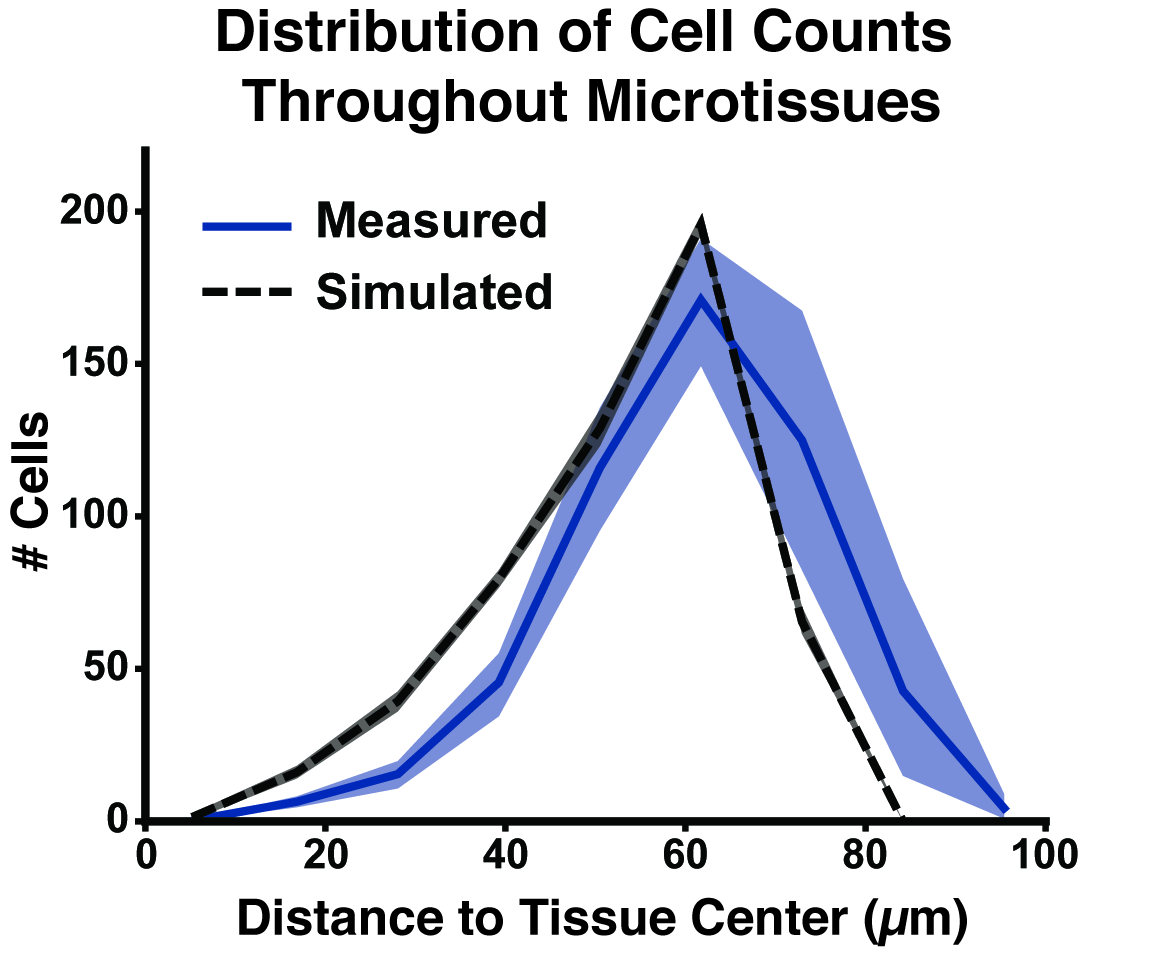

### Supplemental Figure 4

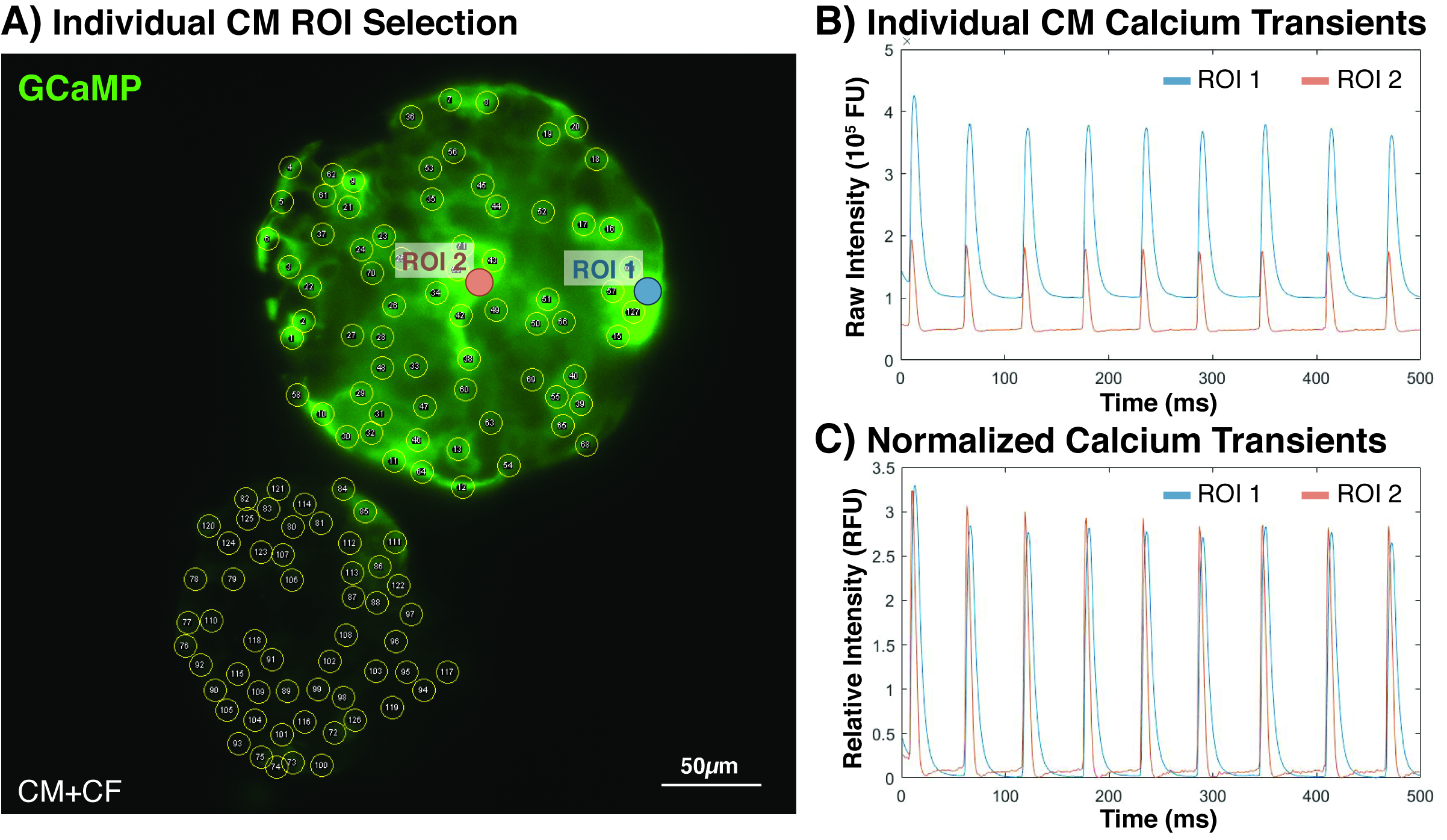
