## Supplemental Table 1 for "Single cell determination of cardiac microtissue structure and function using light sheet microscopy"

**Supplementary Table 1.** Wholemount immunostaining antibody information.

| **Antibody** | **Company** | **Catalog #** | **Dilution** |
| --- | --- | --- | --- |
| GATA4 | Santa Cruz Biotechnology | sc-25310 | 1:50 |
| Alexa Fluor 555 | Thermo Fisher | A-31572 | 1:400 |
| Hoechst | Thermo Fisher | 62249 | 1:1000 |
